# Sub-second heat inactivation of coronavirus

**DOI:** 10.1101/2020.10.05.327528

**Authors:** Yuqian Jiang, Han Zhang, Jose A. Wippold, Jyotsana Gupta, Jing Dai, Paul de Figueiredo, Julian L. Leibowitz, Arum Han

## Abstract

Heat treatment denatures viral proteins that comprise the virion, making virus incapable of infecting a host. Coronavirus (CoV) virions contain single-stranded RNA genomes with a lipid envelope and 4 proteins, 3 of which are associated with the lipid envelope and thus are thought to be easily denatured by heat or surfactant-type chemicals. Prior studies have shown that a temperature of as low as 75 °C and treatment duration of 15 min can effectively inactivate CoV. The applicability of a CoV heat inactivation method greatly depends on the length of time of a heat treatment and the temperature needed to inactivate the virus. With the goal of finding conditions where sub-second heat exposure of CoV can sufficiently inactivate CoV, we designed and developed a simple system that can measure sub-second heat inactivation of CoV. The system is composed of capillary stainless-steel tubing immersed in a temperature-controlled oil bath followed by an ice bath, through which virus solution can be flowed at various speeds. Flowing virus solution at different speeds, along with a real-time temperature monitoring system, allows the virus to be accurately exposed to a desired temperature for various durations of time. Using mouse hepatitis virus (MHV), a beta-coronavirus, as a model system, we identified that 85.2 °C for 0.48 s exposure is sufficient to obtain > 5 Log_10_ reduction in viral titer (starting titer: 5 × 10^7^ PFU/mL), and that when exposed to 83.4 °C for 0.95 s, the virus was completely inactivated (zero titer, > 6 Log_10_ reduction).

**IMPORTANCE:** Three coronaviruses (CoVs) have now caused global outbreaks within the past 20 years, with the COVID19 pandemic caused by SARS-CoV-2 still ongoing. Methods that can rapidly inactivate viruses, especially CoVs, can play critical roles in ensuring public safety and safeguarding personal health. Heat treatment of viruses to inactive them can be an efficient and inexpensive method, with the potential to be incorporated into various human-occupied spaces. In this work, a simple system that can heat-treat viruses for extremely short period was developed and utilized to show that sub-second exposure of CoV to heat is sufficient to inactivate CoV. This opens up the possibility of developing instruments and methods of disinfecting CoV in diverse settings, including rapid liquid disinfection and airborne virus disinfection. The developed method can also be broadly utilized to assess heat sensitivity of viruses other viral pathogens of interest and develop sub-second rapid heat inactivation approaches.

## 1 Introduction

Severe acute respiratory syndrome coronavirus 2 (SARS-CoV-2) is the virus responsible for the currently ongoing global pandemic, the coronavirus disease of 2019 (COVID-19) (1–3). The main transmission routes of SARS-CoV-2 include direct or indirect contact with objects or contaminated surfaces, short-range person-to-person transmission via droplets from coughing or sneezing, and long-range airborne transmission via aerosols (4). Therefore, environmental sterilization and virus inactivation are of great importance to prevent and control the spread of the virus. Currently, the most commonly used methods to sterilize or inactivate viruses include treatment using chemical agents, UV irradiation exposure, and heat treatment, which have all been intensively assessed and reported (5–10). Compared with other methods, one major advantage of heat treatment is its relatively shorter treatment time and simplistic method, along with the ability to be incorporated into various human-occupied space (11–13), which allows for the technique to be readily implemented into a variety of existing applications or systems that could be readily retrofitted to add rapid pathogen inactivation functionality, such as existing heating, ventilation, and air conditioning (HVAC) systems as well as sewer systems.

Heat inactivation is a relatively easy, safe, and efficient method to disinfect coronavirus (CoV), as CoV is an enveloped virus that is surrounded by a lipid bilayer with viral spike proteins projecting from the lipid envelope, where both the envelope and the spike protein are susceptible to heat (14). Previous studies have shown that at a temperature of 56 °C and higher, with heat application time typically longer than 1 min, is needed to efficiently inactivate CoVs such as SARS-CoV and MERS-CoV (> 6 Log_10_ reduction) (5, 6, 13). More specifically, at relatively low treatment temperatures (56 - 65 °C), treatment time of 15 - 60 min was required, while at higher treatment temperatures (70 - 100 °C) a much shorter duration of 1 to 15 min was needed (5, 6, 13, 15–17). For example, heat treatment of SARS-CoV-2 at 70 °C for 5 min achieved > 4.5 Log_10_ reduction (15), with another study reported that heat treatment at 92 °C for 15 min achieved > 6 Log_10_ reduction for SARS-CoV-2 (5). However, for heat treatment to be effectively utilized for liquid and airborne CoV inactivation in broad ranges of practical settings, such methods need to be effective at a significantly shorter heat treatment time (even if the temperature itself has to be higher), otherwise there is limited practicality in such heat treatment methods. For example, having to increase the temperature of liquid for minutes would consume large amount of energy, and having to treat air for minutes becomes impractical.

Here, we hypothesize that a much shorter heat treatment time may be sufficient to destroy key components of CoVs (e.g., envelope or the spike proteins) to completely inactivate CoV. Conventional heat treatment testing method mostly utilizes a simple method of dipping a CoV-containing tube into a temperature-controlled water bath. Such methods are valid when heat treatment time in the range of minutes are utilized, but cannot be used if seconds or sub-second temperature exposure needs to be tested. In this study, we developed a simple flow-through heating and cooling method utilizing a stainless steel capillary tube, and used the method to investigate the effect of CoV heat treatment at an extremely short heat exposure time of 0.1 - 1 s at an actually applied temperature range of 35-100 °C. This study provides essential data for the development of sub-second CoV heat inactivation approaches, including methods to efficiently inactivate airborne CoVs indoors. As all CoVs are surrounded by lipid bilayer membranes with similar proteins and have similar physical properties (14), we expect that our finding using mouse hepatitis virus (MHV), a beta-coronavirus, as a model system, can be broadly applicable to CoVs in general, including SARS-CoV, SARS-CoV-2, and MERS-CoV, to name a few. Since many other viruses, such as dengue virus, influenza virus, and measles virus, are also enveloped viruses where the envelopes contain surface proteins (18), we expect that this heat inactivation method can have broad utility for many other viruses of global consequences.

## 2 Material and Methods

### 2.1 Virus and cells preparation

The coronavirus used in this study is mouse hepatitis virus, strain A59 (MHV-A59), and has been described previously (19, 20). Mouse L2 cells, which are susceptible to MHV infection, were grown in Dulbecco Modified Eagle’s Medium (DMEM) supplemented with 4 mM glutamine and 10% defined calf serum (Hyclone), at 37 °C 5% CO_2_ environment (21).

### 2.2 Virus titration and plaque assay

Plaque assays were conducted as described previously and read at 2 days post-infection after removing the agarose overlay and staining with crystal violet (22). Samples were assayed in triplicate and the plaques were counted, and infectivity was determined and expressed as plaque forming units (PFU)/ml.

### 2.3 Heat treatment of virus solution

Fig. 1 shows the experimental setup that allowed rapid and high-temperature heat treatment of viruses. Stainless-steel (SS) capillary tubing (SS 304, 1/16” outer diameter, 0.02” (500 μm) wall thickness, 0.0225” (572 μm) inner diameter, Mcmaster-Carr, Douglasville, USA) was bent and immersed in an oil bath and then in an ice bath sequentially. A relatively small inner diameter tubing was selected to minimize the volume of solution so that the virus solution can be rapidly heated and cooled down, and SS was utilized to maximize heat conduction from the exterior to the interior of the tubing. Since a short pulse of high-temperature application was desired to accurately assess the impact of the temperature on viral infectivity, an ice bath was utilized to rapidly cool down the heated viral solution. A temperature-controlled oil bath (Instatherm^®^ Economy Bath/Controller Kit, Ace Glass, Inc., VWR, USA) equipped with a type J thermocouple temperature sensor was employed to control the heat treatment temperature. Vegetable cooking oil (Member’s mark, Sam’s Club, College Station, USA) was used in the oil bath. A mixture of ice and water was used as the ice bath to cool down the treated virus solution immediately after treatment. A syringe pump (Fusion 200, Chemyx, Stafford, USA) was employed to control the flow rate of the injected virus solution. Silicon tubing was used to connect to the syringe to the inlet of the SS tubing and also to collect the heat-treated samples from the outlet of the SS tubing. Low-thermal-conducting silicon tubing was utilized to further limit the temperature exposure to only a short section of the overall tubing. The collected virus samples were stored at −80°C until plaque assays were conducted to measure the infectivity of the heat-treated viruses.

**Fig. 1.**
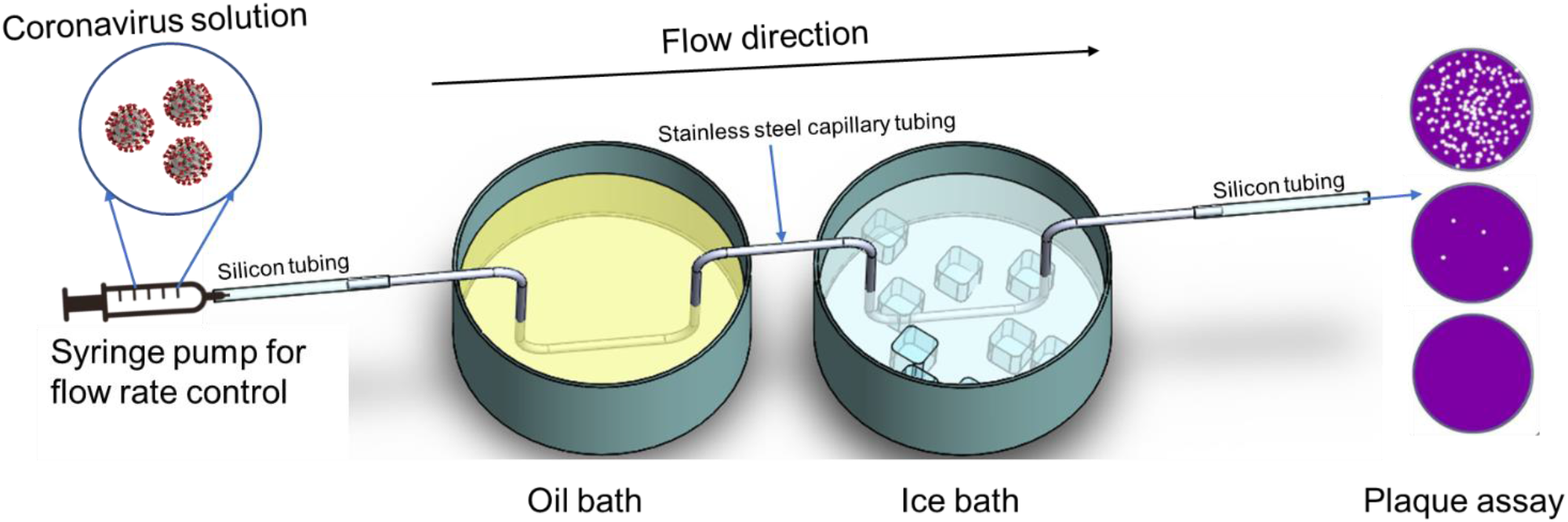
Schematic illustration of the virus heat inactivation system

Stock MHV viruses of 2.7 × 10^9^ PFU/mL were diluted by DMEM media with 10% FBS to 5 × 10^7^ PFU/mL, a high-enough titer to accurately determine the effectiveness of heat treatment on viral infectivity, prior to heat inactivation treatments. For each heat treatment, a virus solution aliquot of 1.5 mL containing approximately 75 million viral PFUs was injected into the SS capillary tubing under varying flow rates by a syringe pump while the oil bath temperature was set as 55 °C - 170 °C. The treated samples were collected at the outlet of the silicon tubing. The set exposure time is calculated based on the traveling distance of the viral solution through the length of the SS tubing immersed in the oil bath (5 cm). The flow rate and set exposure time are summarized in Table S1. A viral solution flowing through the tubing while the oil bath temperature was set to room temperature (22 °C) was used as a control to account for any potential viral titer reduction due to virus adhering to the tubing surfaces and other potential losses.

### 2.4 COMSOL simulation of temperature

The COMSOL Multiphysics 5.5a software (COMSOL Inc., Palo Alto, CA, USA) was used for the temperature profile simulation of the viral solution. To perform this finite element analysis, a 3D geometry was chosen, where the geometry was created using the following initial and boundary conditions: initial temperature of the entire system is set to 20 °C; no-slip conditions on the tubing surfaces; no viscous stress and convective flux on the tubing outlet; the fluid is Newtonian and the flow within the channel is incompressible; convective heat flux is considered as the source of heat influx from the oil to the tubing.

The simulation was performed using non-isothermal flow (nitf) multi-physical interfaces (laminar flow (spf) coupling with heat transfer in solids and fluids (ht) under the stationary study model. The inlet flow temperature and ambient air temperature were set to 22 °C. The material physical properties of viral solution were set to be identical to those of water. The material properties of the SS tubing were set to: density = 7850 kg/m^3^; thermal conductivity = 16.2 W/(m·K); heat capacity at constant pressure = 500 J/(kg·K). Here, we assume the overall heat loss by the oil bath is equal to the overall heat gain of water flowing by (the heat loss to ambient air is neglectable). Then, the temperature-dependent heat transfer coefficient (H) was calculated based on equation (1) (23, 24) and the real-time temperature measurement (see section 2.5 for more details). The calculated H value of 473.6 W/(m^2^·K) for 125°C, 0.5 s and 675.5 W/(m^2^·K) for 170 °C, 0.1 s are close to previously published results (25). Physics-controlled mesh with the element size of “finer” was applied for the simulation.

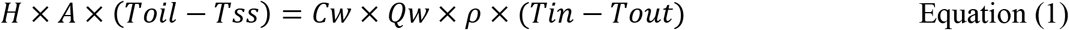

A: surface area where the heat transfer takes place;

T_oil_: temperature of the surrounding oil;

T_ss_: temperature of the solid surface;

C_w_: heat capacity of water;

Q_w_: flow rate of water;

ρ: density of water;

T_in_: water temperature at inlet;

T_out_: water temperature at outlet.

Where,

### 2.5 Real-time temperature measurement and validation of the measurement method

To test the real-time temperature of the viral solution flowing through the tubing while the heat is applied through the heated oil bath, the SS tubing was cut apart between the oil bath and the ice bath, and the two ends were re-connected through a T-connector with insulation material (Fiberglass) wrapped around the connector (Mcmaster-carr, Douglasville, USA). A J-type thermocouple was inserted through the top end of the T-connector to obtain real-time temperature of the viral solution flowing through the tubing. The temperature of the tubing wall was measured by attaching the sensor head of the thermocouple to the outside wall the SS tubing and wrapped with fiberglass.

The temperature measured by this experimental setup (T-thermocouple) was validated and calibrated by withdrawing water heated to a known temperature via a syringe pump (EGATO^®^ 111, KD scientific, MA, USA) while the temperature reading of the sensor was recorded. The schematic illustration of this measurement setup is described in Fig. S1-A.

## 3 Results and Discussion

### 3.1 Validation of the viral solution treatment temperature

As our study aims to inactivate CoVs with a sub-second exposure to high temperature, accurately measuring the real temperature applied to the viral solution is imperative. Since the solution cannot be heated up or cooled down instantaneously and the desired temperature exposure time is quite short, this is non-trivial. First, to validate whether the thermocouple inserted into the T-junction of the SS tube (called T-thermocouple here) can accurately measure the solution temperature inside the SS tube, a solution with known temperature, set by heating the bulk of the solution in a temperature-controlled water bath, was drawn into the SS tubing at various flow speeds and its temperature was measured. This measured temperature inside the SS tubing (2 cm away from the heated water bath) was then compared to that of the bulk solution temperature. Fig. S1-B in the ESI shows these calibration curves. The results show that the temperature probe accurately measures the solution temperature inside the SS tube. For example, at a fast flow of 0.1 s exposure, which minimizes any cooling of the solution outside of the water bath, the measured temperature was almost identical to the water bath temperature (Fig. S1-B, 0.1 s condition, 97.8 °C real temperature = 97.7 °C measured temperature). Second, as the flow rate becomes slower (*i.e.*, longer exposure to heat), it is expected that the solution will cool down slightly the moment it leaves the SS tubing section immersed inside the water bath. Indeed, the measurement result shows that as the exposure time increases to 0.25, 0.5, and 1 s, the measured temperature becomes slightly lower than the water bath temperature. For example, for water bath temperature of 98 °C and exposure time of 0.25, 0.5, and 1 s, the T-thermocouple temperature reading was 95.4, 94.5, and 92.6 °C, showing the slight cooling of the temperature. Therefore, the real-time measured temperatures were corrected by the calibration curves (Fig. S1-B), which were used as the actual temperatures applied.

### 3.2 COMSOL simulation

The heat transfer efficiency and temperature profile within the SS capillary tubing are important to understand the actual heating condition of the viral solution. We used COMSOL Multiphysics^TM^ to simulate the heat transfer from the SS tubing to the flowing CoV solution inside the tubing. Fig. 2 (A) shows the temperature change of flowing virus solution within the SS tubing when the oil bath temperature was set to 125 °C and a moderate exposure time of 0.5 s (flow rate = 92300 μl/hr) was used. Here, the SS tubing temperature increases rapidly as it enters the oil bath (pink zone), then gradually drops when the tubing is exposed to air (yellow zone), and then rapidly drops as it enters the ice bath (blue zone). The reason the SS temperature inside the oil bath (pink zone) is not same as the oil bath temperature is due to the cooling effect of room temperature water continuously flowing into the tubing, especially since the length of the tubing immersed inside the oil bath was relatively short (5 cm). Using a much longer tubing inside the oil bath would have eventually made the SS tubing temperature same as the oil bath temperature, but that makes it impossible to apply a short temperature exposure time, hence the use of a relatively short tubing length.

**Fig. 2.**
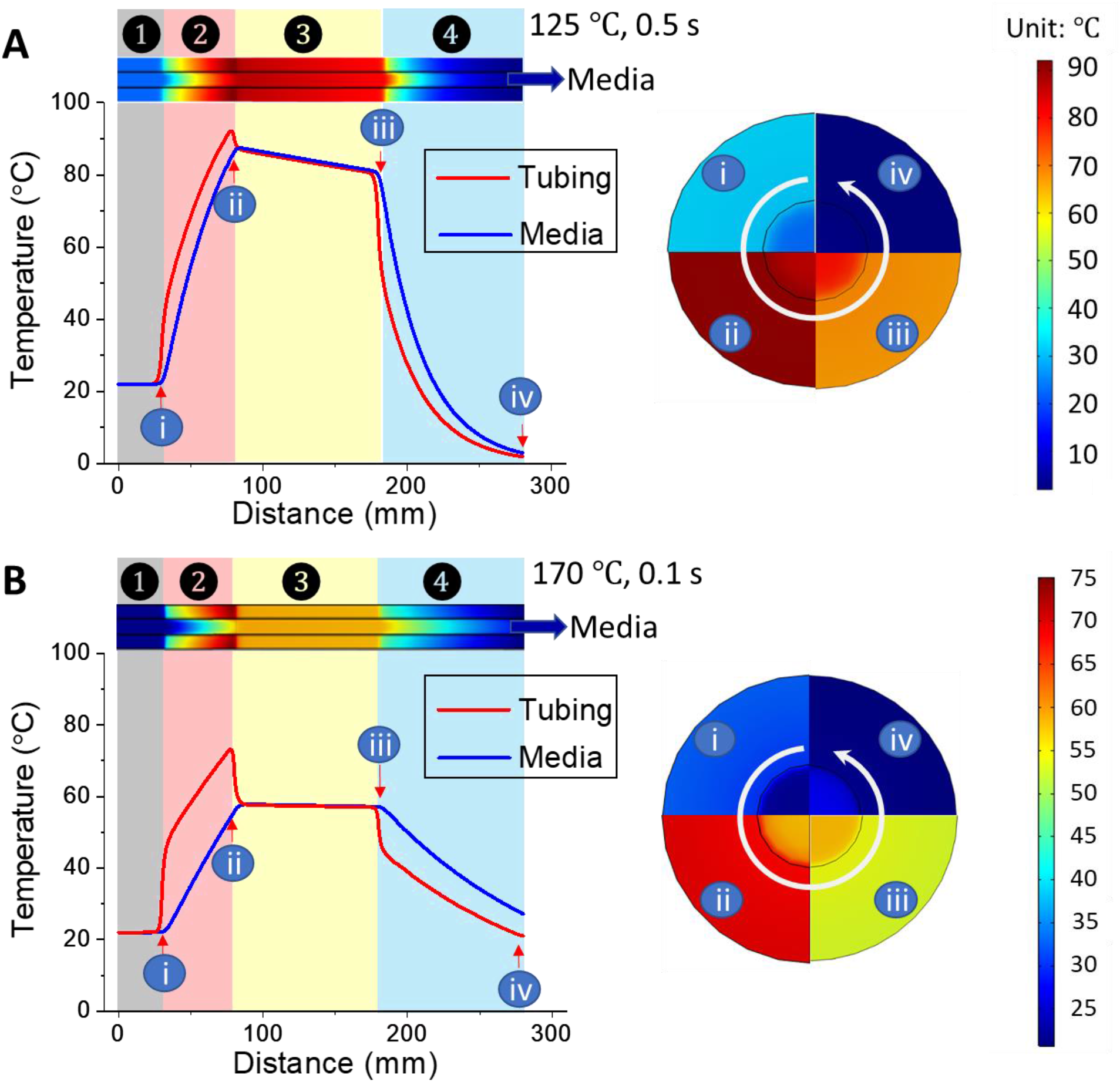
The simulated temperature distribution of the entire heat inactivation system when using oil bath to apply and sectional views of positions i - iv are displayed in the right circle: (A) 125 °C, 0.5 s exposure, and (B) 170 °C, 0.1 s exposure. Zone ① pre-oil bath; Zone ② oil bath; Zone ③ ambient air; Zone ④ Ice bath.

The solution temperature follows the SS tubing temperature relatively closely. The highest solution temperature of 87 °C was achieved right as it comes out from the oil bath and maintains its temperature relatively well (temperature drop of 6.5 °C over a 10 cm length), until arriving at the segment of the SS tubing immersed in the ice bath. Therefore, the effective heat treatment region was designated from the highest temperature point post-oil bath and right before the ice bath treatment, a length of 9.5 cm. While the “applied exposure time” is calculated based on the tubing length immersed in the oil bath (5.0 cm), the simulated actual exposure time is 1.9 times longer than the set time, in this case 0.95 s (when the applied treatment time was set to 0.5 s).

Fig. 2 (B) exhibits the temperature distribution under the condition of 170 °C oil bath temperature and “set exposure time” of 0.1 s (flow rate = 461500 μl/h). Even though the set temperature was higher, due to five times faster flow rate of incoming room temperature solution, the SS tubing temperature only reached 56 °C due to the larger cooling effect coming from the incoming solution. This effect can also be seen by the drop in the SS tubing temperature as it exits out of the oil bath. As in the previous case, the highest solution temperature of 56 °C was at the point where the SS tubing exits the oil bath, with minimum drop in temperature until it enters the ice bath. Thus, as before, the effective heat treatment region could be designated from the highest temperature point post-oil bath and right before the ice bath treatment, a length of ~ 9.5 cm. The temperature comparison table summarizes (Table S1, ESI) the oil bath set temperature, the real-time T-junction temperature sensor readout of the CoV solution, the calibrated CoV solution temperature to accommodate the cooling effect, and the simulated temperature.

### 3.3 Heat inactivation of MHV

As reported, an average viral load of SARS-CoV-2 is 7 × 10^6^ per milliliter (26), therefore we chose 5 × 10^7^ PFU/mL of MHV which is slightly higher and would provide a rigorous test of this rapid heat inactivation system. Our study investigated rapid heat treatment, within one second, to demonstrate effective coronavirus inactivation, by flowing MHV virus solution through stainless tubing immersed in an oil with temperatures ranging from 55 °C to 170 °C. We found that the coronavirus particles were inactivated efficiently (> 6 Log_10_ reduction) by applying an oil bath temperature of 115 °C and residence time for 1 s. When the heat exposure time decreased to 0.5 s, a temperature of 125 °C or higher is needed to completely inactivate the MHV. When the heat exposure time further decreased to 0.25 s, the virus titer reduction is 5 Log_10_ at 150 °C treatment. However, when the heat exposure time reduced to 0.1 s, the MHV in the capillary stainless-steel could not be sufficiently inactivated even when the oil bath temperature was increased to 170 °C.

Therefore, the remaining infectivity after heat treatments comparing different conditions was re-plotted by the actual thermal treatment temperature and duration, and is shown in Fig. 3. Among the set oil bath temperatures ranging from 55 °C to 170 °C, the shortest treatment time required to inactivate the MHV effectively (> 5 Log_10_ reduction) was concluded as 0.25 s (0.48 s as simulated), with an actual exposure temperature of 85.2 °C. Furthermore, the most rapid thermal treatment to completely inactivate the MHV (> 6 Log_10_ reduction and the remaining titer is 0 PFU/mL) is 0.5 s (0.95 s as simulated), with an actual exposure temperature of 83.4 °C.

**Fig. 3.**
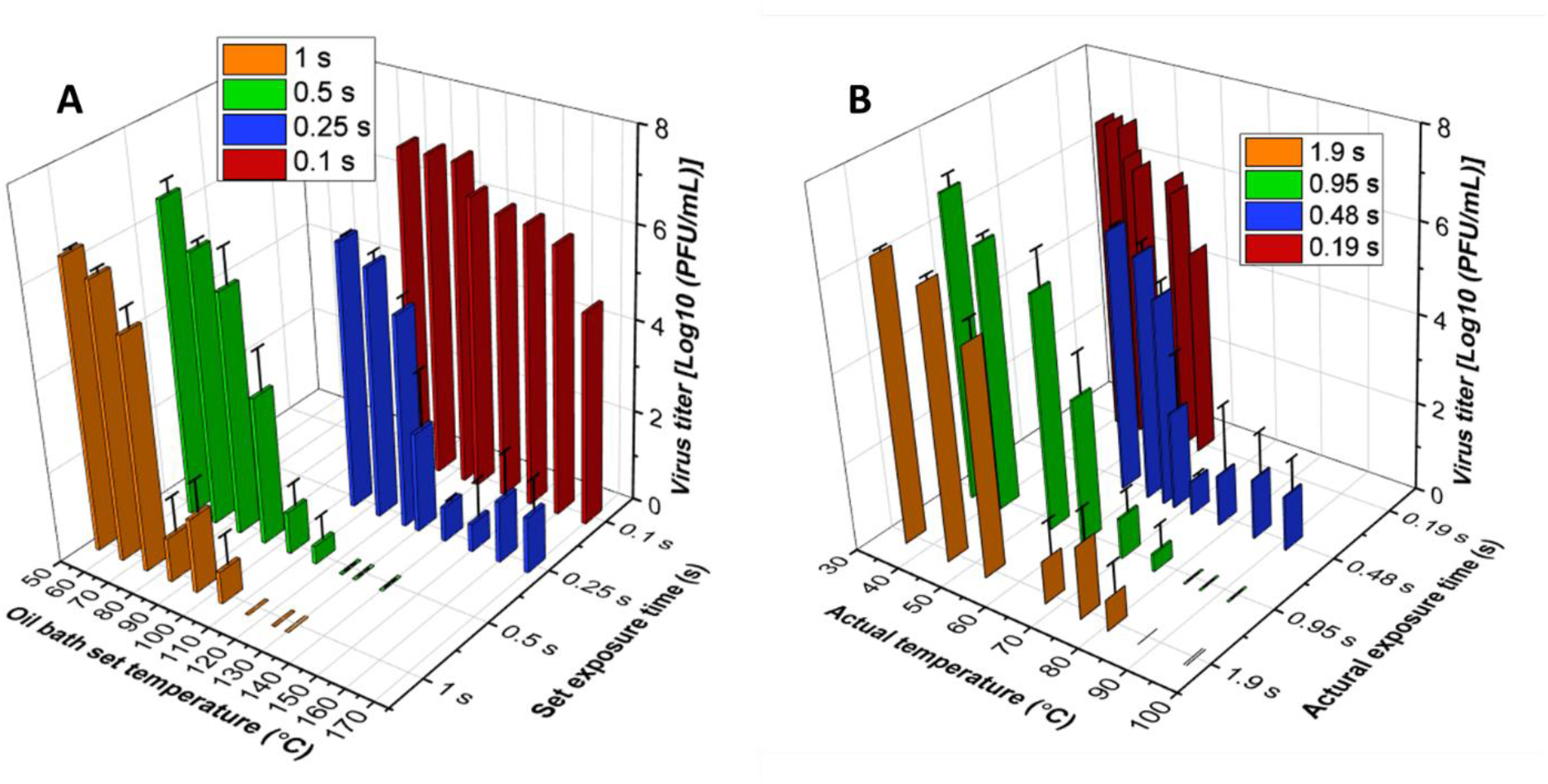
Effect of heat treatment on the infectivity of MHV. Remaining infectivity of MHV after different heat inactivation conditions (A) by oil bath temperature and set exposure time; (B) by actual temperature and exposure time from real-time measurement. Virus titers were averaged from three independent biological replicates (n = 3), and error bars indicate standard deviations.

Table 1 summarizes the inactivation effectiveness under different heat treatment conditions, where our results were shaded. Generally, coronaviruses are relatively stable at room temperature and are sensitive to heat when the temperature increased to 65 °C, after which the sensitivity destruction linked to temperature became obvious. As the small increases in temperature (after 65 °C) cause a large increase in death rate, and temperature experienced by the viruses would not equal the applied heating temperature (27), our study fills the gap of sub-second heat treatment under applied actual temperature.

**Table 1.**
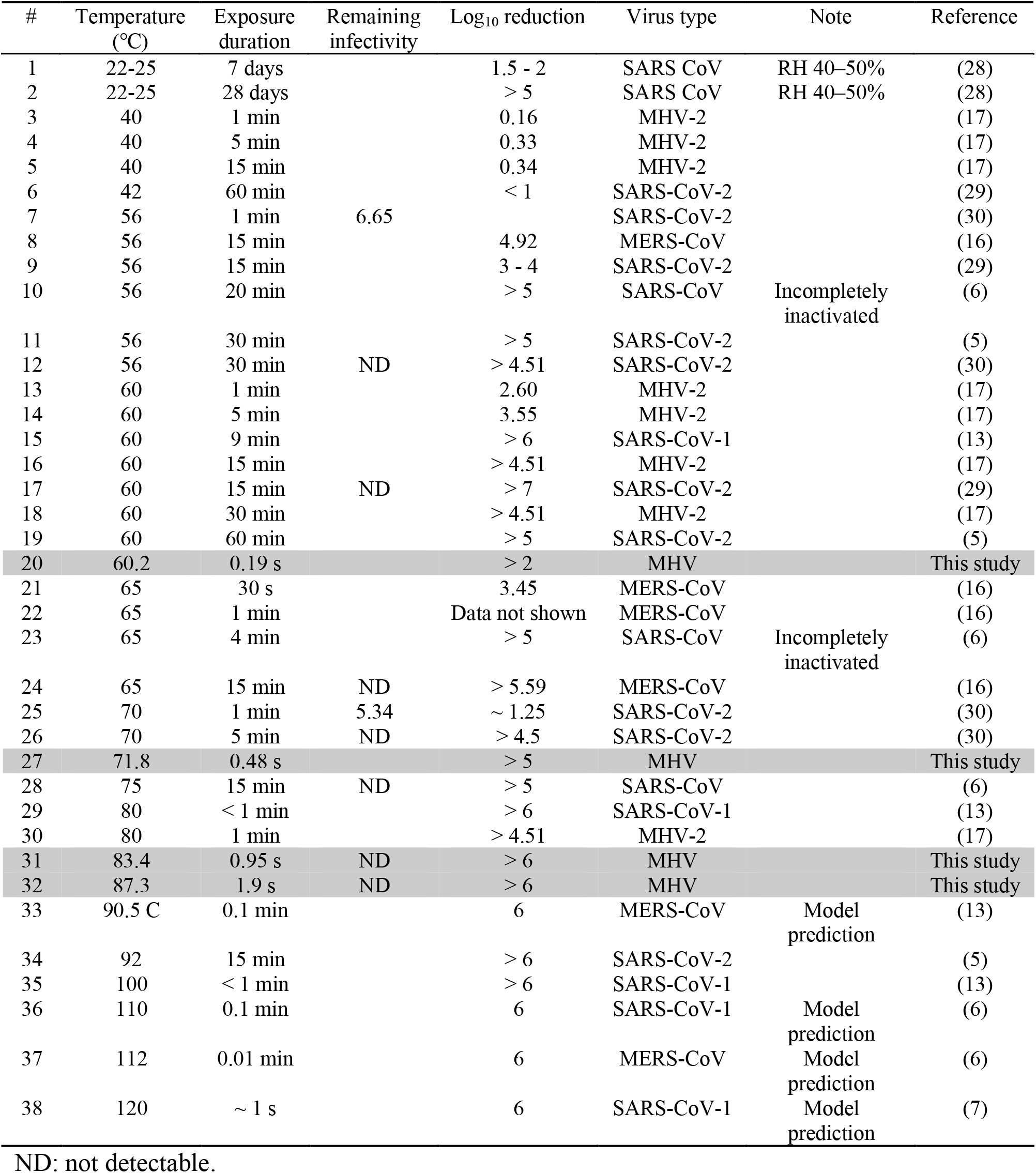
Summary of reported heat inactivation conditions with temperature ranging from room temperature to 120 °C for heating duration among sub-second to 1 h.

## 4 Conclusion

We studied the thermal treatment conditions to efficiently inactivate a coronavirus and developed a system and protocol that can inactivate infectibility of MHV. We successfully developed an experimental setup that allows applying sub-second duration of heat to the CoV solution where the temperature applied to the CoV solution can be monitored in real time. Through experimental measurement and computational thermal simulation, we validated the real temperature that the CoV solutions are exposed to. Using this setup, we identified that 85.2 °C for 0.48 s exposure is sufficient to obtain > 5 Log_10_ reduction in viral titer (starting titer: 5 × 10^7^ PFU/mL), and that when exposed to 83.4 °C for 0.95 s, the virus was completely inactivated (zero titer, > 6 Log_10_ reduction). This is the first experimental result that shows that sub-second exposure to high temperature is sufficient to inactive the infectibility of CoV. Since heat treatment is a simple, inexpensive, and efficient approach to inactivate coronaviruses, our method can be used to study the thermal sensitivity of viruses, as well as providing critical data that can be used to develop efficient CoV heat inactivation methods that can be broadly applied to real-world settings.

## Acknowledgements

This project was supported by Medistar Corporation. We would like to thank Dr. Garrett K. Peel (MD, MHS, FACS) and Mr. Monzer Hourani from Medistar Corporation for their valuable discussion and input throughout the project. We would like to also thank Dr. Victor Ugaz who provided invaluable insights into the simulation aspect of the project. Some research personnel on this project were also partially supported by the National Institute of Allergy and Infectious Diseases (NIAID) grant 1R01AI141607-01A1.

